# A heat method to interpolate z-stacked cell images that preserves interfaces without explicit identification of object boundaries

**DOI:** 10.1101/2025.04.11.648268

**Authors:** Donald L. Elbert

**Author notes:** Corresponding author: Donald L. Elbert.

## Abstract

A multitude of methods have been developed to morph one image into another. In scientific and medical imaging, it is common to have lower resolution in one imaging direction. To improve the quality of surface meshes generated from these images, morphing is used to produce intermediate images to achieve equal resolution in all directions. The most common method is based on interpolating signed distance functions that describe the shapes of objects in the images. Most methods rely on identification and recording of object boundaries. The situation becomes more complex in the case of interpolating images containing multiple objects of interest wherein the morphing method must not introduce gaps or overlaps of objects in the intermediate images. The goal of the present work was to develop an interpolation method for biological cells in transmission electron microscopy images meeting the constraints listed above. A heat diffusion method was developed that produces steady state isotherms in a 3D model of the intervening space between two images. The isotherms are then used to produce paths from pixels that are unique to a biological cell in one frame towards the overlap region. Instead of identifying boundaries, only membership in the overlap region is tested. For each pixel that terminates paths, the path lengths of all members are normalized to the maximum path length. The path lengths are then used to produce intermediate frames. The method succeeds in avoiding the introduction of gaps or overlap between neighboring biological cells without the need for boundary identification.

## Introduction

As part of a larger project, we are generating smoothed 3D surface meshes of cells in the brain. In a companion paper, we describe the nature of the problem in detail.^1,2^ In brief, the data are serial sectioned TEM images of mouse visual cortex.^3^ The segmented images contain some errors that are not consistent with membrane biophysics. As will be shown here through examples, the most severe segmentation errors seem to be associated with the algorithm used to interpolate missing frames in the TEM data, which is performed by a convolutional neural network used for alignment and segmentation.^4,5^ Additionally, the resolution in the x and y directions are 5 times higher than in the z-direction, which will affect the quality of surface meshes that are generated from the data. Appropriate smoothing and interpolation schemes are vital to further progress on the overall project.

Interpolation of frames in a z-stack of images is a common issue in the generation of 3D objects from medical imaging data.^6,7^ It is desirable to generate cubic voxels rather than rectangular voxels to improve the quality of 3D meshes.^7^ If images are segmented manually or automatically, each object may be defined by its signed distance function.^6,7^ Adjacent images represented by their signed distance functions may be linearly or cubic spline interpolated, with a new boundary drawn at the zero crossing.^7–9^ While this works well for single objects, the current problem involves hundreds of objects with defined interfaces that must be respected by not introducing overlaps or gaps. As an example, where one cell branches, cells residing in the saddle region must maintain contact with each other and the branching cell. How the cells in the saddle region fill the space is undefined from the images so one of many plausible solutions must be chosen in some way. Using differences in signed distance functions, it would be difficult to avoid overlaps and gaps between the cells in the interpolated frames.

Classical image morphing techniques that are used to blend source and target images seek to produce smooth transitions by introducing intermediate frames primarily as a video effect. The simplest technique is to simply adjust the transparency of pixels to ‘cross-dissolve’ or linearly interpolate between the two images.^10^ This leads to an unacceptable artifact of ‘double exposure’ or ‘ghosting’ in intermediate frames that is unconvincing due to the lack of respect for the boundaries of the source and target objects. A groundbreaking improvement was introduced by Industrial Light & Magic called ‘mesh warping’.^11^ This involves the identification of landmarks such as facial features to produce corresponding meshes in both images. The source mesh is then warped into the target mesh, with a cross-dissolve within the warped mesh.^11^ The ‘deform and cross-dissolve’ method was applied to medical images by Ruprecht and Müller.^12,13^ Landmark identification was originally performed manually. Sederburg and Greenwood began to address the automation problem by considering morphing of one polygon into another for cases without topological changes, which eventually led to an automated morphing method.^14,15^ Mesh or landmark based interpolation could be applied to the current problem, but the presence of tens to hundreds of objects with meandering boundaries and topology changes between frames complicate any procedure that relies on boundary processing.

For example, one solution would be to identify cell boundaries in both images, develop meshes within the non-overlapping regions of the cell in each image and solve the associated heat transfer problem, with one boundary being a Dirichlet boundary condition and the other an insulating Neumann boundary condition. This would produce appropriate morphing of cells but involves a great deal of boundary bookkeeping and mesh generation. Herein is presented a method that does not require identification of cell boundaries and solves the heat transfer problem on a uniform Cartesian mesh.

## Results

The thin sections imaged by transmission electron microscopy are nominally 40 nm in thickness. The resolution of the TEM images is 4 nm and segmentation voxels are 8 nm x 8 nm x 40 nm. Thus the x and y-directions have a resolution that is five times finer than in the z-direction. If Marching Cubes is applied to the raw segmentation images, the resulting meshes are stair-stepped and elongated in the z-direction (Figure 1A). Elongated (skewed) elements lead to loss of accuracy in computational methods.^16^ Downsampling in the x and y directions is feasible, but there are structures (such as dendritic spines) that are sometimes not much thicker than 40 nm (Figure 1B). Another possibility is to repeat each layer five times, again leading to stair stepped meshes but with high quality elements (Figure 1C). In addition to stair stepping, there are spikes and ledges that are very thin (8 nm). Such structures are too small to be part of the plasma membrane. As will be shown, these are almost certainly the result of segmentation errors. A series of filtering and morphing steps will be described in the manuscript that produce high resolution meshes with high quality elements (Figure 1D).

**Figure 1:**
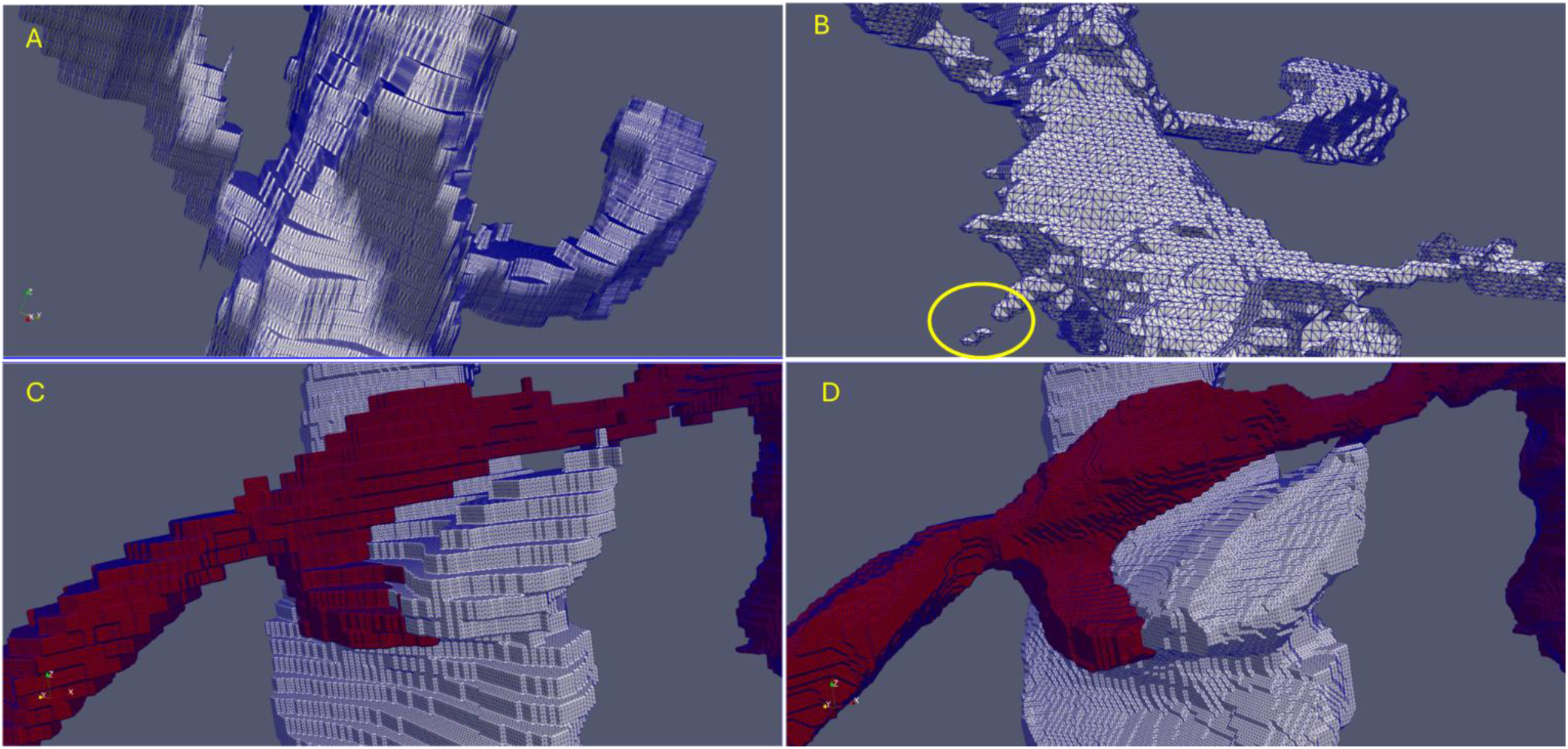
A. The mismatch between the resolution in the x and y directions (8 nm) and the z direction (40 nm) may be addressed by stretching elements in the z-direction, but this decreases the quality of the mesh. B. Alternatively, downsampling the voxels in the x and y directions may lead to fragmentation of small dendritic spines (yellow circle). C. Another option is to repeat each z layer five times, producing high quality elements but a stair-stepped mesh. D. Finally, the new mode filtering and image morphing method described here produces smoother meshes with high quality elements.

Zooming in on Figure 1C, the source of the ‘ledge’ and ‘spike’ are illustrated in Figure 2. The likely source of these structures is due to a single missing frame, in which the interpolation algorithm assigns spurious pixels to a cell. The interpolation is not an algorithm per se, but rather the result of training a convolutional neural net with missing frames.^4^ The raw images do not support the presence of the ledge and spike (data not shown but may be examined at MicronsExplorer cortical-mm3 site at location x=340583, y=122407, z=17415-17; https://www.microns-explorer.org/cortical-mm3).

**Figure 2:**
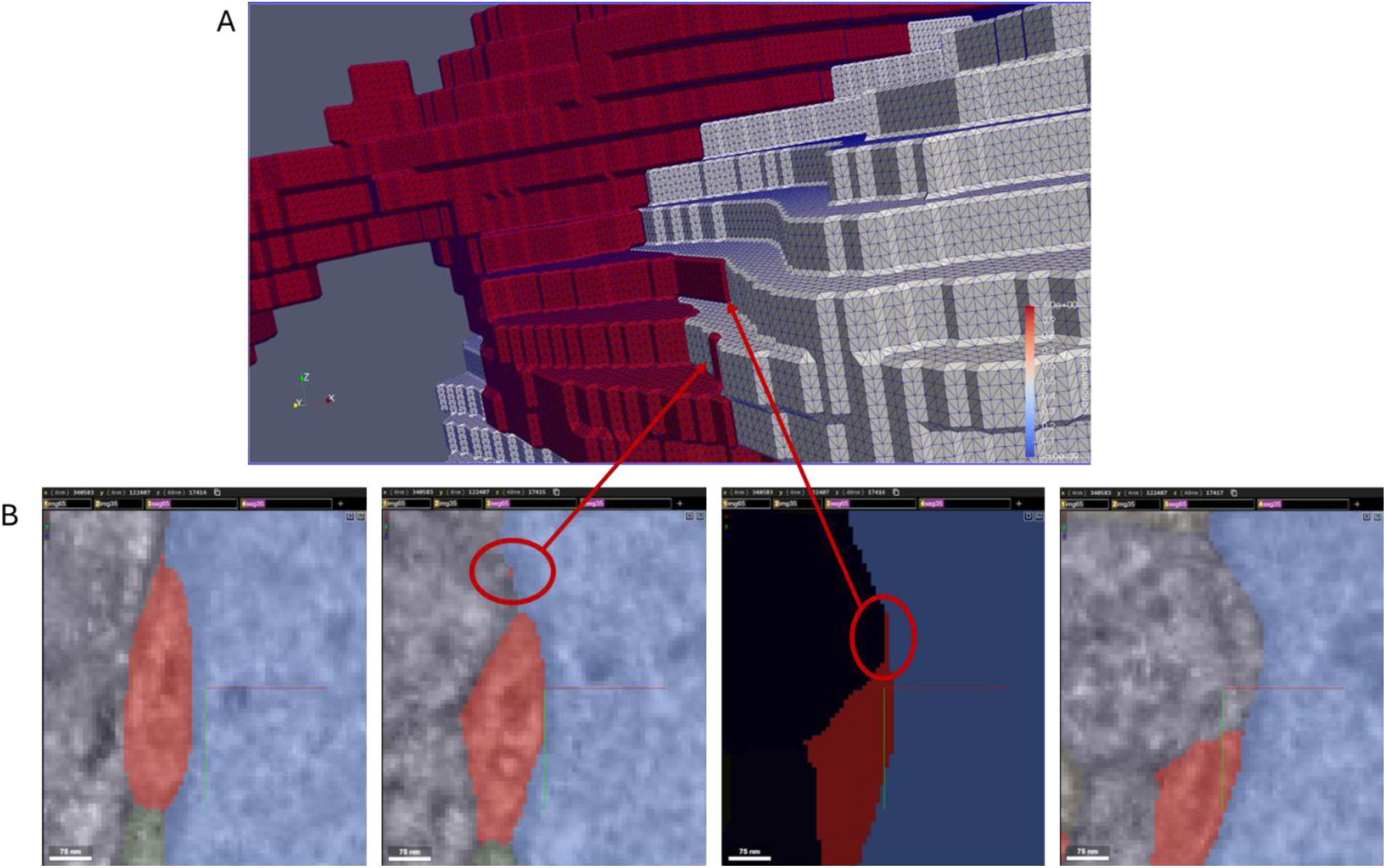
Illustrations of segmentation artifacts that result in thin prominences and thin ledges. The implied diameter/width of these structures is 8 nm, which is not biophysically plausible. These structures are often adjacent to missing frames and may result from the image morphing algorithm used to fill in missing frames. Scale bars are 75 nm.

Additional spikes and other prominences present in the segmented images are shown in Figure 3A&B. The yellow circled base of two spikes is due to isolated pixels assigned to cell 864691135666883810 in neighboring frames (using the v343 nomenclature). Once again the TEM images (not shown) do not support the presence of these spikes. These spikes and an adjacent ledge result in a highly twisted surface mesh, with a 180-degree rotation of face normals for faces attached to the same vertex (Figure 3A). Figure 3B shows another case, where a convergence of four cells 864691135128790164, 864691136697855485, 864691135478307526 (not shown) and 864691136328907498 (not shown) and single pixel protrusion from cell ‘164’ in the frame below (inset) results in a hole in the surface mesh. This type of structure will severely limit the quality of volumetric meshes that model the extracellular space, which may be produced by adding ‘thickness’ to the surface meshes. After mode filtering and morphing, the twists and holes are no longer present and the surface mesh is smooth (Figure 3C&D).

**Figure 3:**
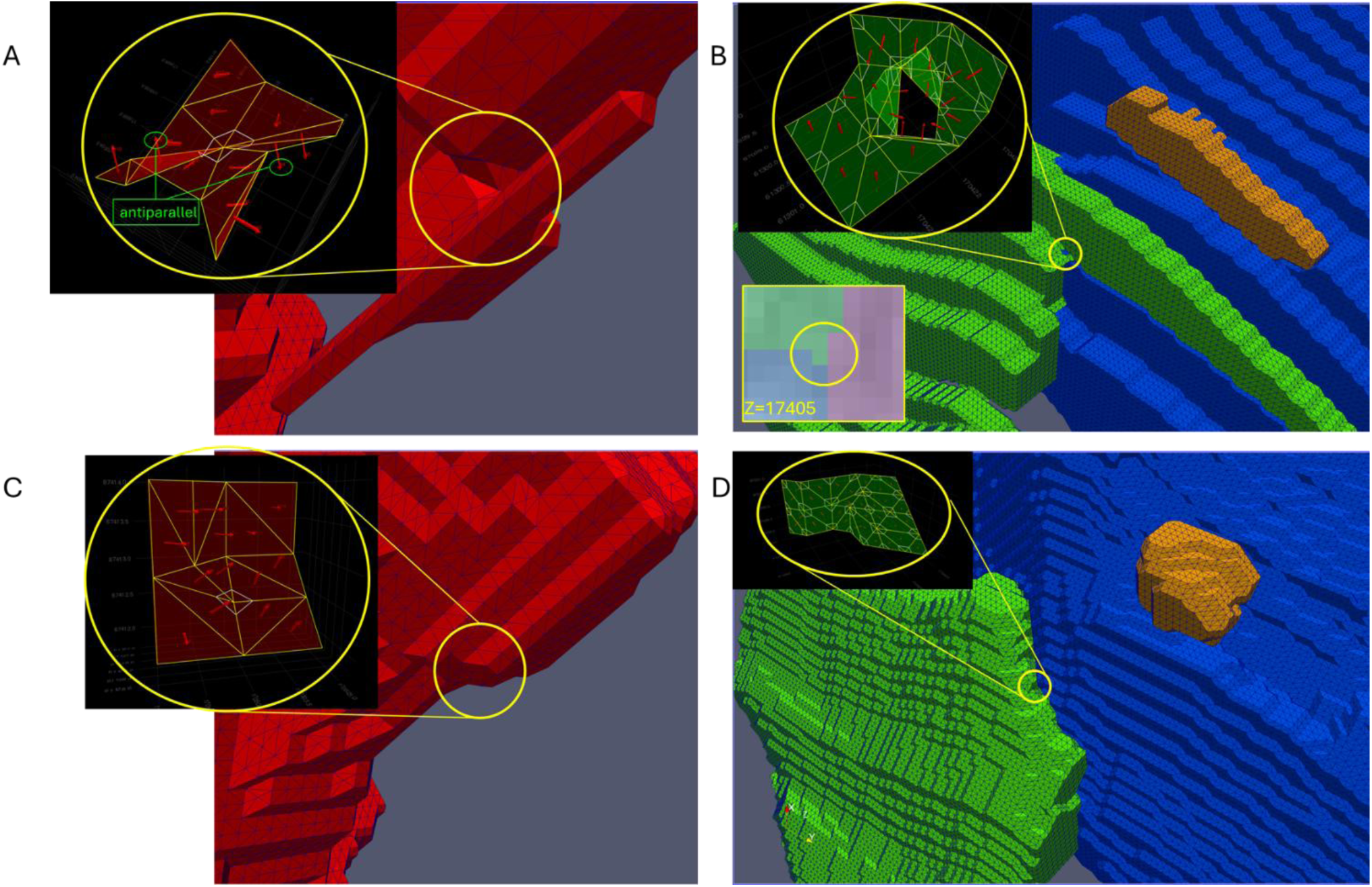
Complex geometries of surface meshes in raw segmented data. A. Due to the presence of single isolated pixels in the frames above and below the circled region, ‘spikes’ result when each layer is repeated five times. A region of the surface mesh in the inset shows that at a single vertex, the surface mesh wraps 180° around the vertex to accommodate the spike. Cell 864691135666883810; x=341010, y=122256, z=17483. B. A surface mesh wraps around a hole. The hole is due to the convergence of four cells and a single pixel protrusion in the lower frame (inset). Green = cell 864691135128790164; x=340844, y=122602, z=17406. Blue = cell 864691136697855485. Orange = cell 864691132493097261. C&D. After smoothing and morphing, simpler surface mesh geometries are found. Panel C corresponds to panel A and panel D corresponds to panel B.

Mode filtering by itself is sufficient to remove spikes and ledges. Figure 4A shows a mesh derived from the unsmoothed segmented images (blue) overlayed with the mesh resulting after mode filtering (green). Focusing on the circled spikes in Figure 4A&B, the isolated pixels assigned to this cell are shown in white in the raw segmented image (Figure 4B). The inset shows the resulting image, with the cell of interest in blue. The isolated pixels are far away from the main body of the cell and once again almost certainly are artifactual. Both of the isolated pixels were removed by a single round of mode filtering. The modified mode filter algorithm is shown in Figure 4C. Figure 4Ci is the traditional mode filter. However, where three or more cells meet, there may be a three-way tie. In this case, the center pixel is marked for removal and is replaced by the cell with lower cell number (Figure 4Cii). There are also cases where the center pixel is the mode but forms a single pixel prominence. In this case, the center pixel is marked for removal even if it is the mode. The rule is applied when the mode count is four and less than two off-diagonal pixels are the same as the center pixel. This will preserve square corners and limit erosion of cells during successive application of the mode filter. However, the rule also preserves some elongated thin shapes that are not desirable but these will be removed by repeated application of the mode filter. The final rule (Figure 4Civ) removes small structures. If a cell is assigned to the center square but the cell is not present on the edge of the surrounding 11 x 11 window, the pixel is marked for removal. If a pixel is marked for removal, it cannot be the mode and thus the center pixel is the second most common entry. There are some objects in the segmentation that are too small to be cells and often appear to be spines that have been separated from their dendrite (e.g. orange ‘cell’ in Figures 3B, which is about 300 nm x 60 nm x 60 nm in size). Handling these cell fragments automatically would obviate the need for the rule in Figure 4iv.

**Figure 4:**
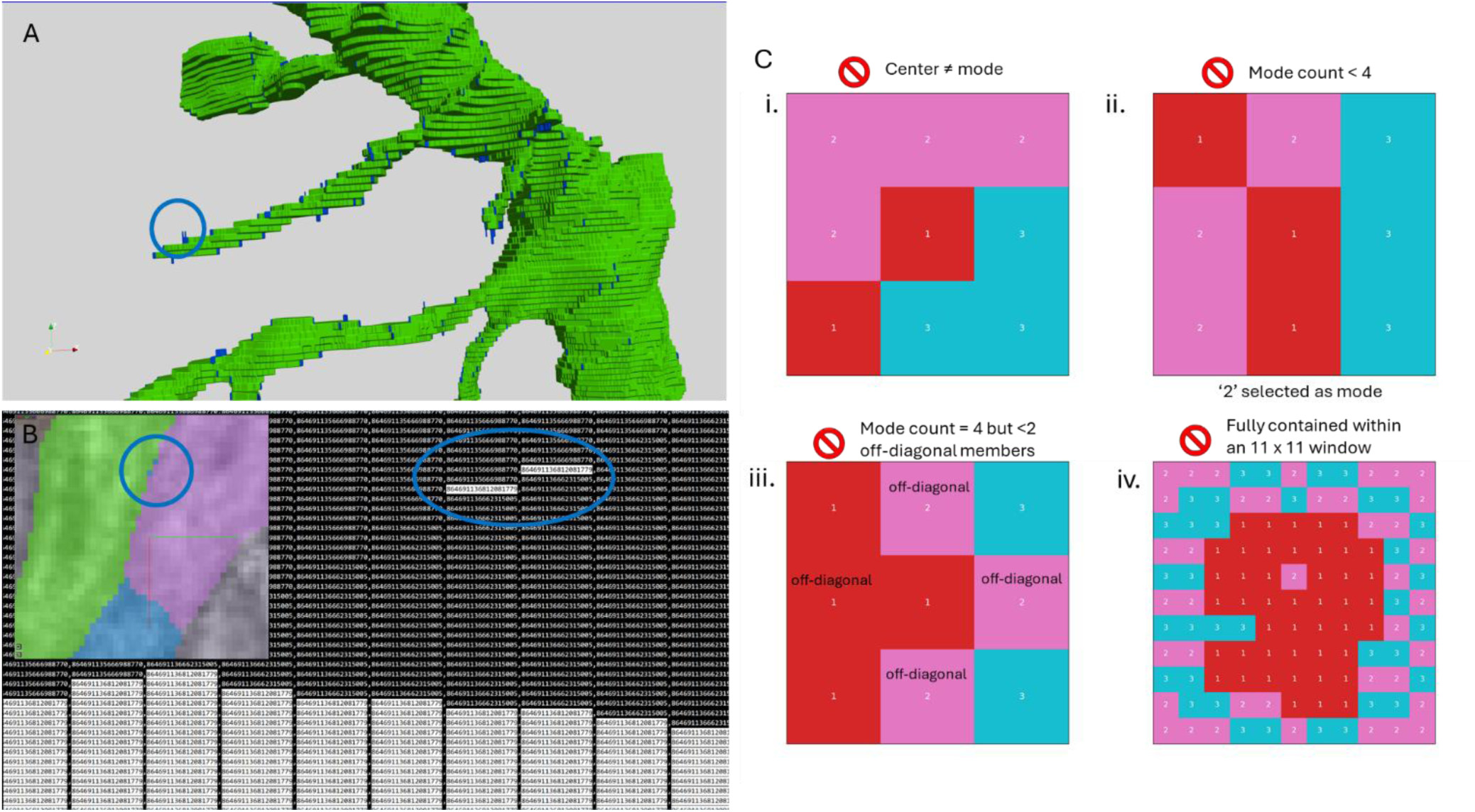
A) Prominences in blue were removed by mode filtering. B) A raw segmented frame showing the eighteen digit cell codes assigned to each pixel. Cell 864691136812081779 is highlighted in white. The circled pixels correspond to the circled prominence in (A). The inset is the false colored image of the segmentated frame with cell ‘779’ shown in blue. C) Description of mode filter: (i) traditional mode filter. (ii-iv) Special rules added to the traditional mode filter were developed for: (ii) interfaces between 3 or more cells, (iii) preservation of square edges, and (iv) removal of isolated segments that were smaller than 11 x 11 pixels.

After mode filtering, adjacent images in the z-stack are interpolated using the new morphing protocol. To achieve a z resolution of 8 nm, four interpolated frames are introduced, each with a nominal thickness of 8 nm. Figure 5 highlights an example that will be used throughout the rest of this document. The cell in light blue pushes the orange and pink cells apart over the course of one frame and then makes contact with the cyan cell. As seen in these images and those in Figure 2, changes between frames are substantial but not so large that the cells become difficult to track between frames.

**Figure 5:**
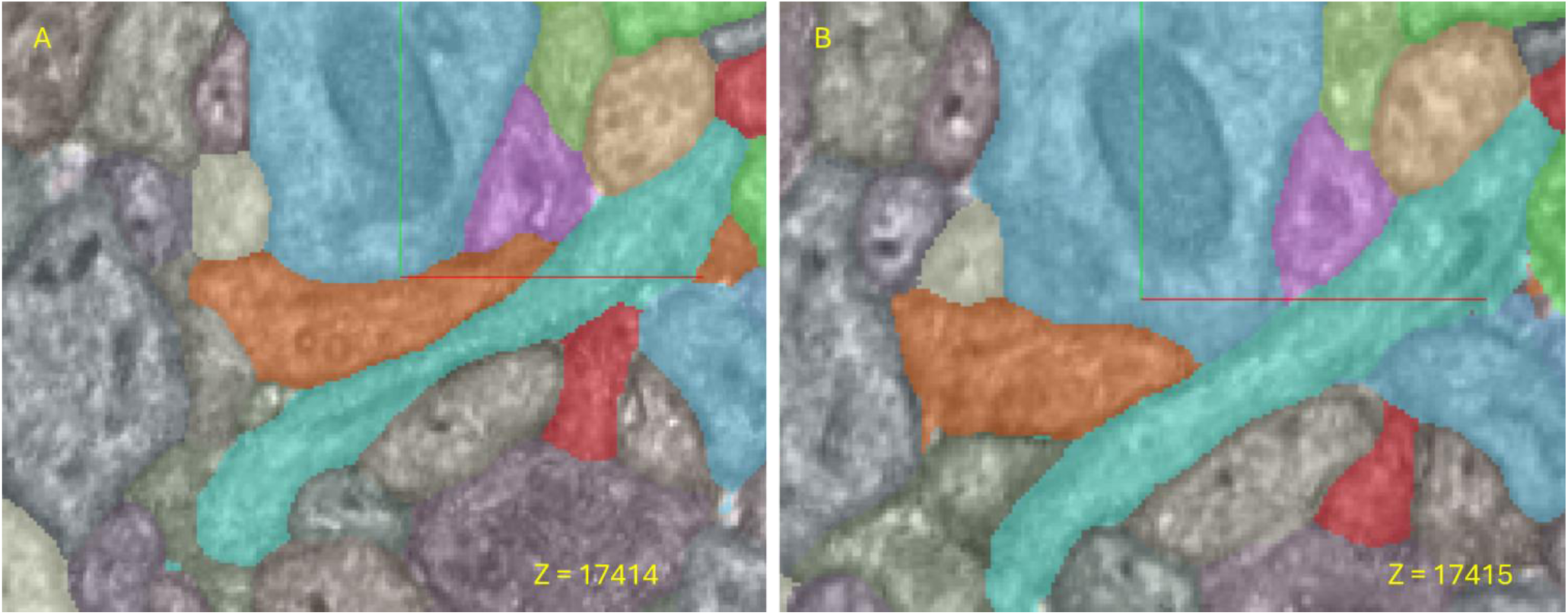
Between frames 17414 and 17415, the orange and pink cells retract and the light blue cell pushes towards the cyan cell. Center x = 340924, center y = 122420 in 4 nm coordinates from the Microns Explorer website (https://www.microns-explorer.org/cortical-mm3). Axis line lengths are about 1 μm.

A biological cell may have more than one domain (i.e. multiple components) within the frames. In Figure 6A, an upper frame (pink) and lower frame (green) of a single distinct biological cell are in an otherwise empty frame, with two distinct components in the lower frame and one component in the upper frame. Intermediate frames will need to capture the branching of the cell in between the two frames. The shape of the branches should minimize membrane energetics in some way, but the exact shape is impossible to deduce from the images. The shape of the branch may vary considerably so any reasonable solution is plausible. While it is tempting to map the cell boundaries from one frame to the other, this is made much more difficult in the case of branching (Figure 6B).^12^

**Figure 6:**
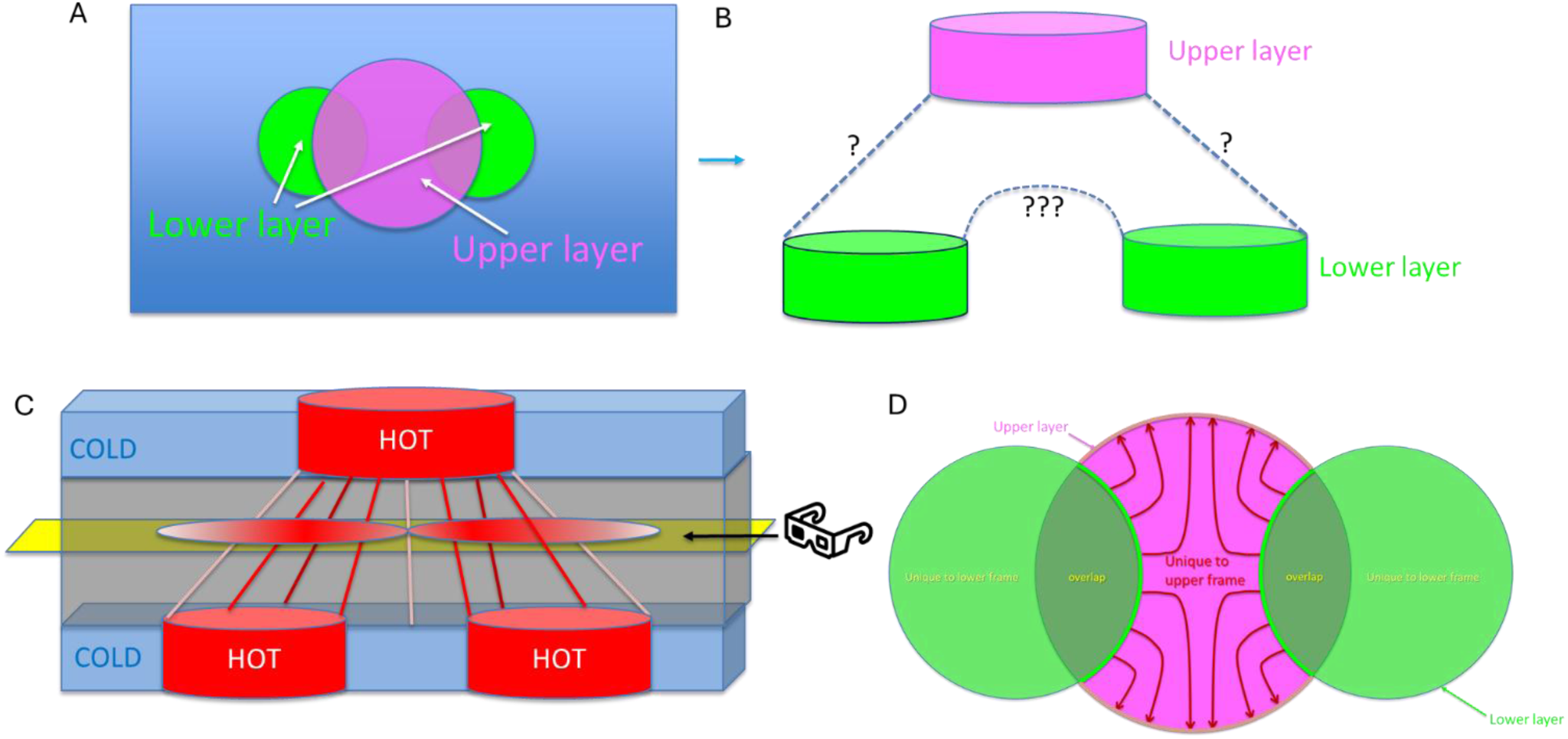
New image morphing protocol illustrated with a branching cell. A. A single cell occupies one region in the upper frame (pink) but occupies two distinct regions in the lower frame (green). B. In 3D this would represent branching, which presents challenges for directly mapping the lower boundary to the upper boundary. C. A new method was developed that does not require identification of boundaries. In the new method, an intervening heat conducting layer is introduced, sandwiched between the frames. In the top and bottom layer, pixels associated with a unique cell are assigned a non-zero temperature while the surrounding pixels are assigned zero temperature. Boundary conditions are applied only at the outer boundary to the volume, allowing use of a uniform Cartesian mesh. The heat conduction equations are solved numerically to steady state separately for each unique cell present in the two frames. Isotherms are recorded in an observation layer (yellow). D. The final step is to define paths between the green border of the overlap region and the pink border of the region unique to the upper frame.

The method that was chosen is a variant of heat diffusion-based morphing techniques but differs by introducing a third dimension to the heat diffusion problem. The solution was to introduce 25 layers between the image frames, which are described by a 3D rectangular grid that conducts heat. The entire frame is modeled but only one distinct biological cell (possibly with multiple components) is present in the pair of frames. The biological cell components are assigned a ‘hot’ value of temperature to the cell (e.g. temp = 1.0) and then ‘imbedded’ in an otherwise cold layer (temp = 0.0). The other boundary faces of the volume are fixed to temp = 0.0. The diffusion equation is discretized by the finite volume method on the cubic grid with Δx, Δy and Δz = 8 nm (or more properly, set to 1 unit, where 1 unit = 8 nm). The solver is run to steady state and the solutions at discrete z-levels are stored for later use. For efficient storage, temperatures less than 0.05 were set to zero and non-zero values were saved in a sparse matrix. The distance between the upper and lower frames (200 nm) does not need to agree with the true distance of 40 nm as only the isotherms at steady state at various levels are of interest.

Observing heat transfer in the intermediate zone (gray), areas of cell overlap will have the highest temperatures, while other areas will have intermediate temperatures combined with a fast drop off in temperature the region occupied by cell components (Figure 6C). The final step is to devise rules for cell movement between layers (Figure 6D). This involves identifying paths from the overlap boundary (thick green border) to the boundary of the region unique to the upper frame (thick pink border). The reason that paths are only in the pink region is because the non-overlap region that is green for this cell is within the pink region of other cells. Collectively, other cells will overrun the green regions of this cell. The choice of region for migration is arbitrary although the resulting morphed images would be slightly different if migration was within the green region instead of the pink region.

In Figure 7, the lower frame is Z = 17414 (Figure 7A) and the upper frame is Z = 17415 (Figure 7B). Unique cells have different colors and the cell in purple will be the focus moving forward. This is the same region as shown in Figure 4, where the light blue cell in that image corresponds to the purple cell in this figure. It is evident that the thick green borders correspond to cell boundaries in the lower frame, while pink borders correspond to the cell boundaries in the upper frame. Figure 7C shows only the green and pink borders, illustrating how they cross each other multiple times per cell (on average) producing a large number of domains that would need to be identified and tracked in most morphing approaches. Using the heat transfer model outlined in Figure 6, the resulting steady state temperature contour lines (isotherms) for each cell is shown in Figure 7D.

**Figure 7:**
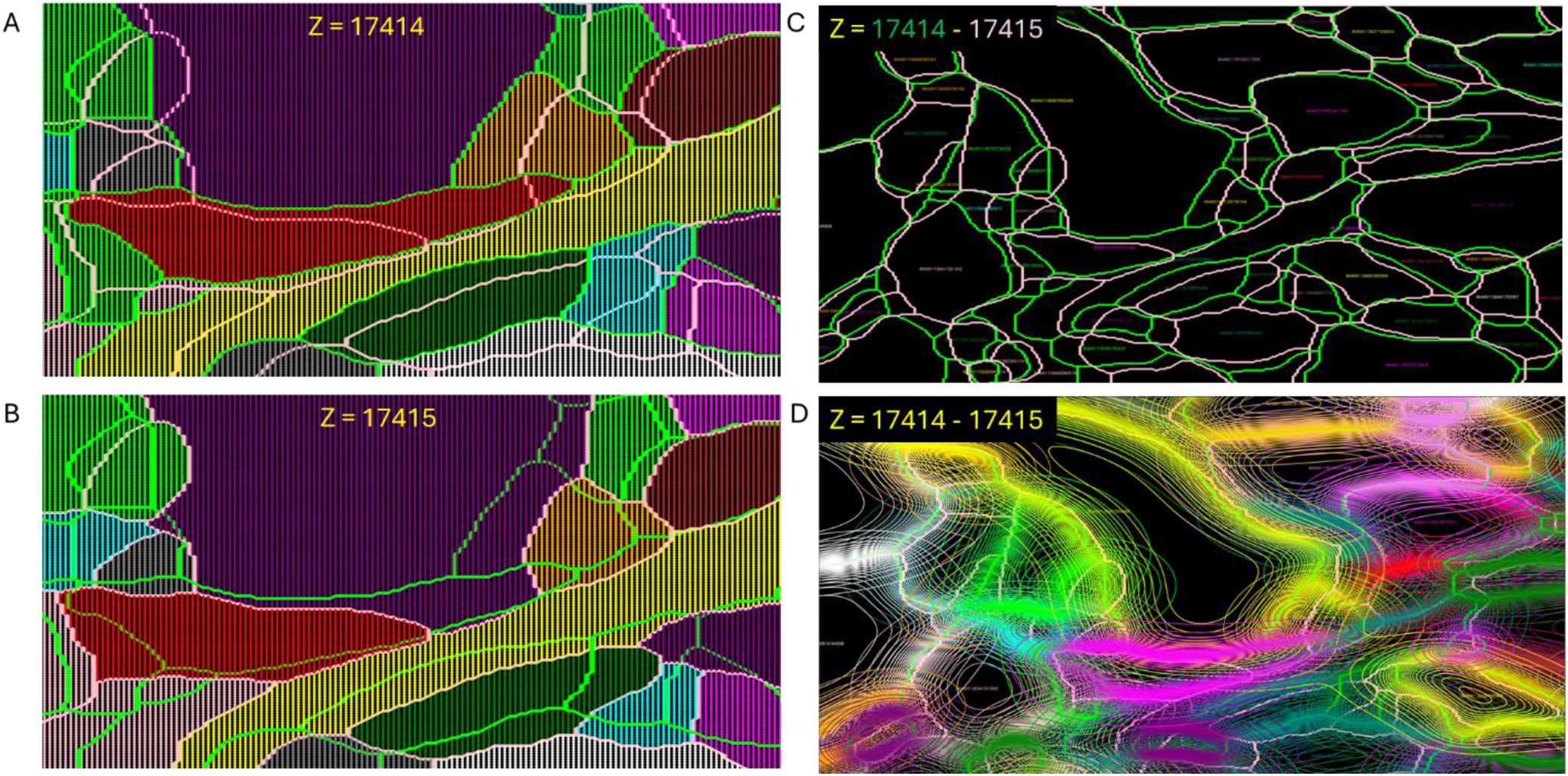
A. Unique cells in the lower frame (17414) are shown in different colors and are contained within the green boundaries. B. Cells in the upper frame (17415) follow the pink boundaries. C. The green and pink boundaries overlayed. D. The contour lines (isotherms) from the steady state solution of the heat equation for each cell.

To determine migration paths, an approach was adopted that is illustrated in Figure 8. The approach was to scan all of the pixels that contained the cell of interest and were unique to the upper frame. For each of these pixels, a path was developed by following the temperature gradient, using gradient ascent with α = 1. At intermediate steps, the path location was usually off of grid points so the gradient at the new location was interpolated from the gradient at surrounding grid points. After each step, the nearest grid point to the current location was checked to determine if it was present in the overlap region (where the cell is in both frames). If so, the path was terminated and the starting pixel coordinates and total path length were stored in a dictionary of dictionaries where the key is a tuple of the x and y coordinates of the terminal pixel. After scanning all of the pixels in the regions that are unique to the upper frame, the dictionary of terminal pixels was analyzed to determine the longest path that reached each terminal pixel. Note that the terminal pixels were almost always on boundaries, but occasionally a path would reach a pixel just beyond the boundary. This would lead to an inaccurate assignment of path lengths at that non-boundary pixel, but it was rare enough that the error would be corrected by subsequent mode filtering. Accepting these occasional errors results in an algorithm that does not require identification of boundary pixels.

**Figure 8:**
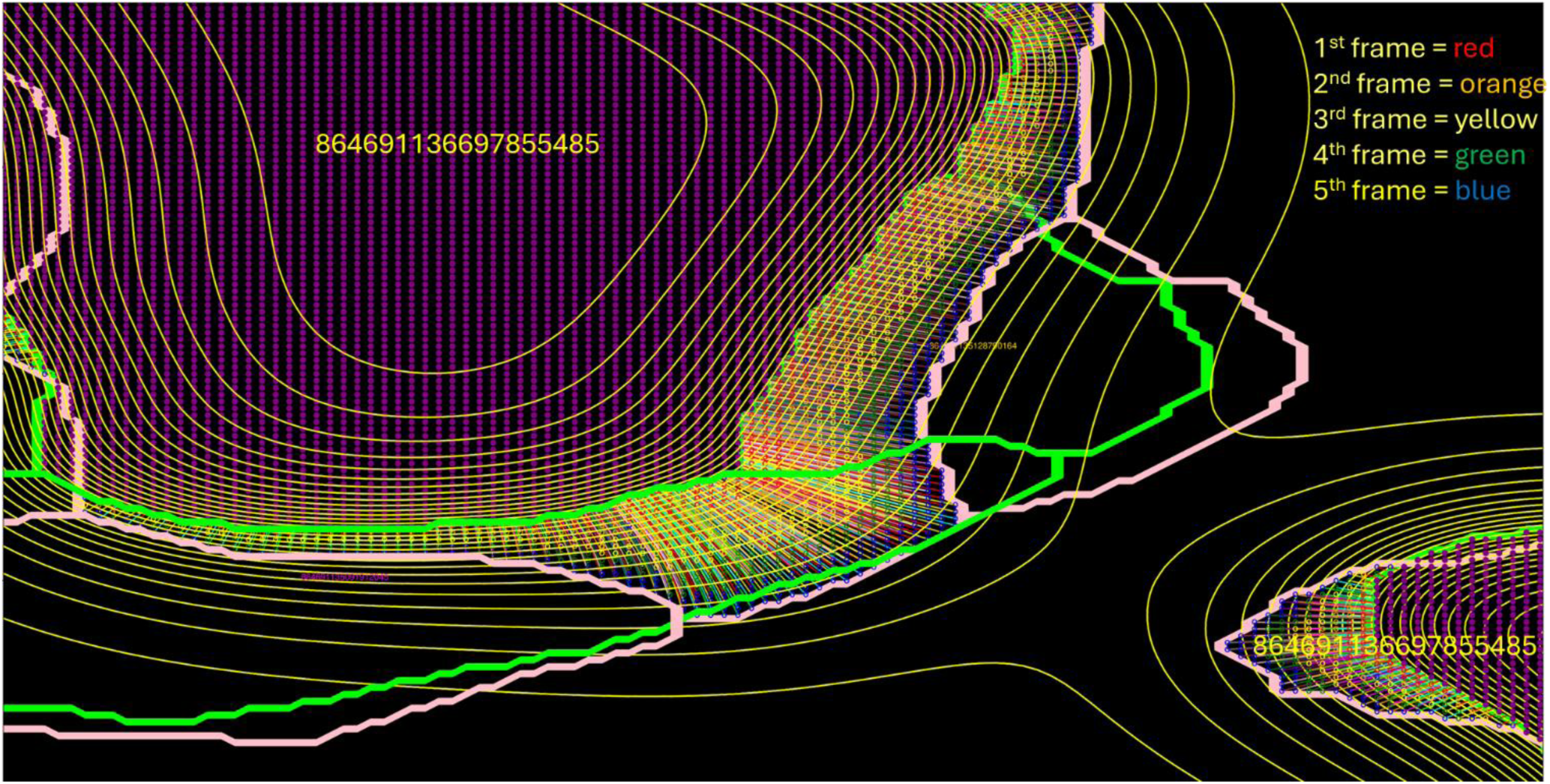
Pixels in the lower frame corresponding to cell 864691136697855485 (‘485’) are shown in purple. ‘485’ is present in two distinct regions (upper left and lower right). The ‘morphing region’ contains pixels that belong to ‘485’ in the upper frame but not in the lower frame. For each pixel in the morphing region, isotherms are followed until the nearest pixel is in both the lower and upper frame (‘overlap region’) and this pixel is stored as an ‘terminal pixel’. The green boundary is not defined explicitly, but instead a list of terminal pixels is maintained. After analysis of all pixels in morphing regions, the list of paths attached to a terminal pixel is iterated, and the longest path associated with the terminal pixel is recorded. The length of all other paths terminating at the same pixel are normalized by the longest path. Paths that are 20% of the longest path define pixels that are converted to cell ‘485’ in the first morphed frame, paths that are 40% of the longest path are converted to ‘485’ in the second morphed frame, etc. Rarely, a path will miss the true boundary and terminate just beyond the green boundary. This is rare enough that mode filtering will correct these errors.

The next step would be to assign a ‘color’ to each pixel in the region unique to the upper frame. For all of the pixels that terminate at the same pixel, those with path lengths <20% of the maximum path length associated with that terminal pixel were assigned a color of red in Figure 8. Path lengths 20<=x<40% of the maximum path length were orange, 40<=x<60% of the maximum path were yellow, 60<=x<80% were green and the remaining pixels were blue.

For the first frame inserted above the lower frame, any pixel colored red is assigned to the cell in its terminal pixel. For the next frame, any pixels colored red or orange are assigned to that same cell. This repeats until four new frames have been generated. The final frame (with blue pixels) does not need to be generated as it is the upper frame. Following production of the interpolated frames, the entire volume is mode filtered again, not only in the xy plane, but also the yz plane, zx plane and finally the xy plane again.

The resulting intermediate frames (after mode filtering) are shown in Figure 9. The frame numbers (i.e. z-locations) are multiplied by five to avoid fractional frame numbers. Circled in purple is the region shown in Figure 8. The transitions are relatively smooth but mode filtering does tend to lead to fast closure of small gaps. Circled in red is a region that disappears between the two original frames. In the morphed images, this disappearance is not only smooth, it also coincides with the smooth inflow of neighboring cells.

**Figure 9:**
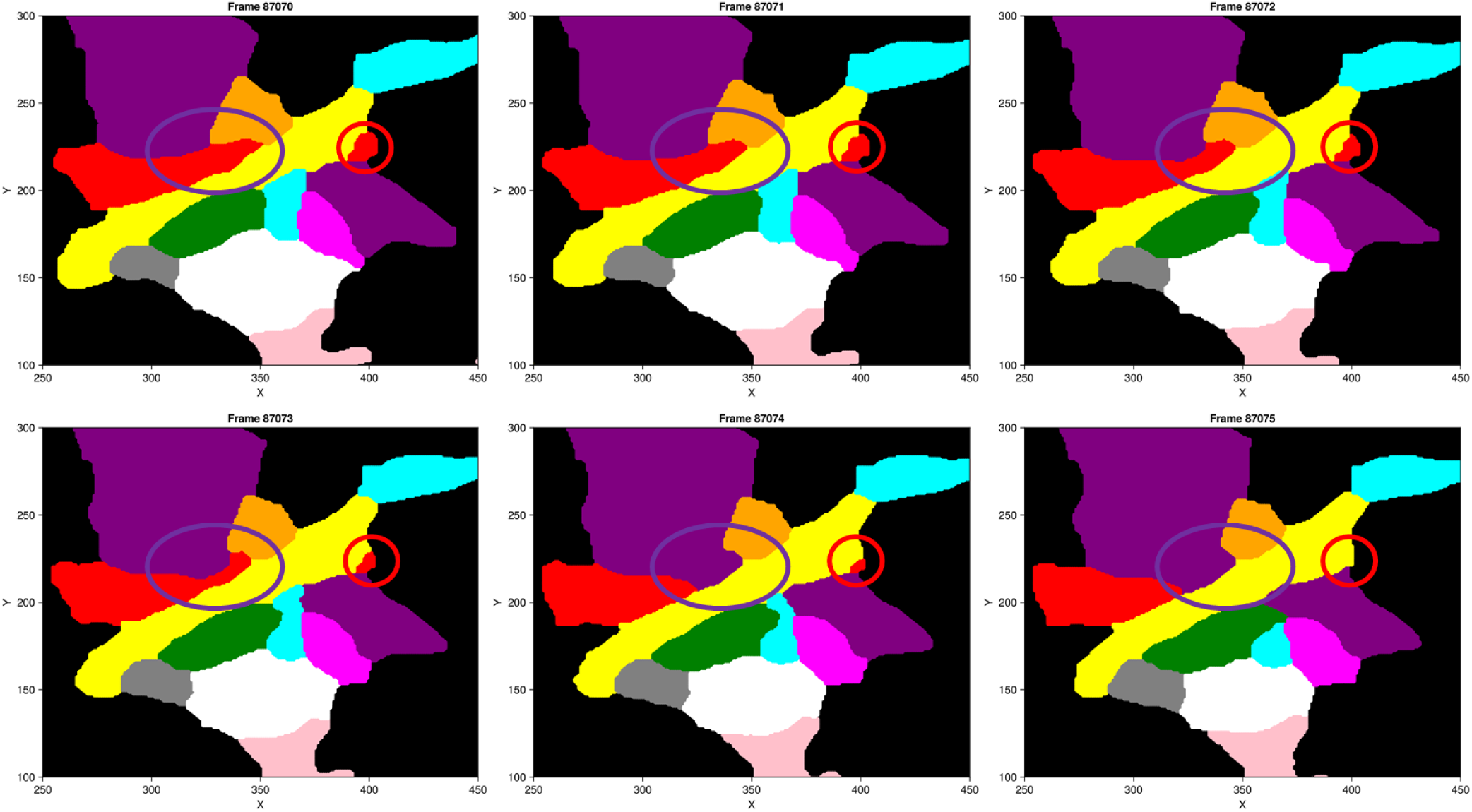
The first (zframe = 87070) and last (zframe = 87075) frames are the mode filtered segmentations. To avoid fractional frame numbers, original frame numbers are multiplied by 5, i.e. zframe 87070 is the same as frame 17414 and zframe 87075 is the same as frame 17415. Intermediate images were generated by the heat diffusion method described in Figure 6, which was followed by additional mode filtering in the xy plane, zy plane xz plane and the xy plane again. Circled in purple, the ‘485’ cell (purple) smoothly pushes away the red and orange cells. Circled in red, part of the red cell disappears between the first and last frame and the transition is shown in the intermediate frames.

Figure 10 shows the next pair of frames. Frame Z = 17415 (Figure 10A) corresponds to Figure 7B, however, the green and pink boundaries are different because here frame Z = 17415 is the lower frame, not the upper frame. Between frame Z = 17415 and frame Z = 17416 (Figure 10B), two domains of the purple cell fuse, corresponding to the challenging case presented in Figure 6. A close-up view of the purple cell is shown in Figure 11, with the paths added to pixels in the region unique to the second frame. There exists a local minimum in temperature (dotted yellow circle).

**Figure 10:**
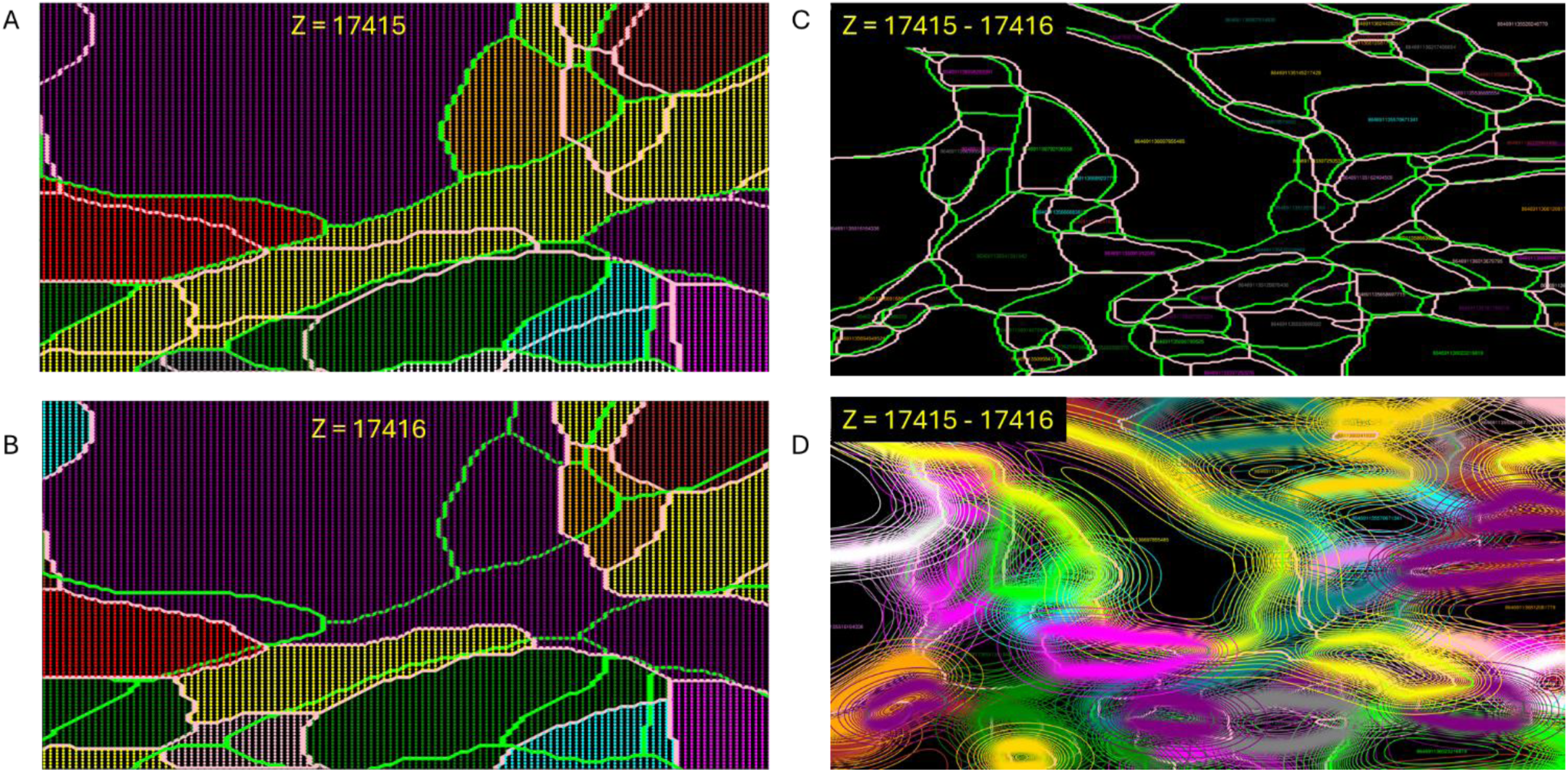
The next pair of frames in the series. A. Unique cells in the lower frame (17415) are shown in different colors and are contained within the green boundaries. B. Cells in the upper frame (17416) follow the pink boundaries. C. The green and pink boundaries overlayed. D. The isotherms from the steady state solution of the heat equation for each cell. Compare panel A here to panel B in Figure 7.

**Figure 11:**
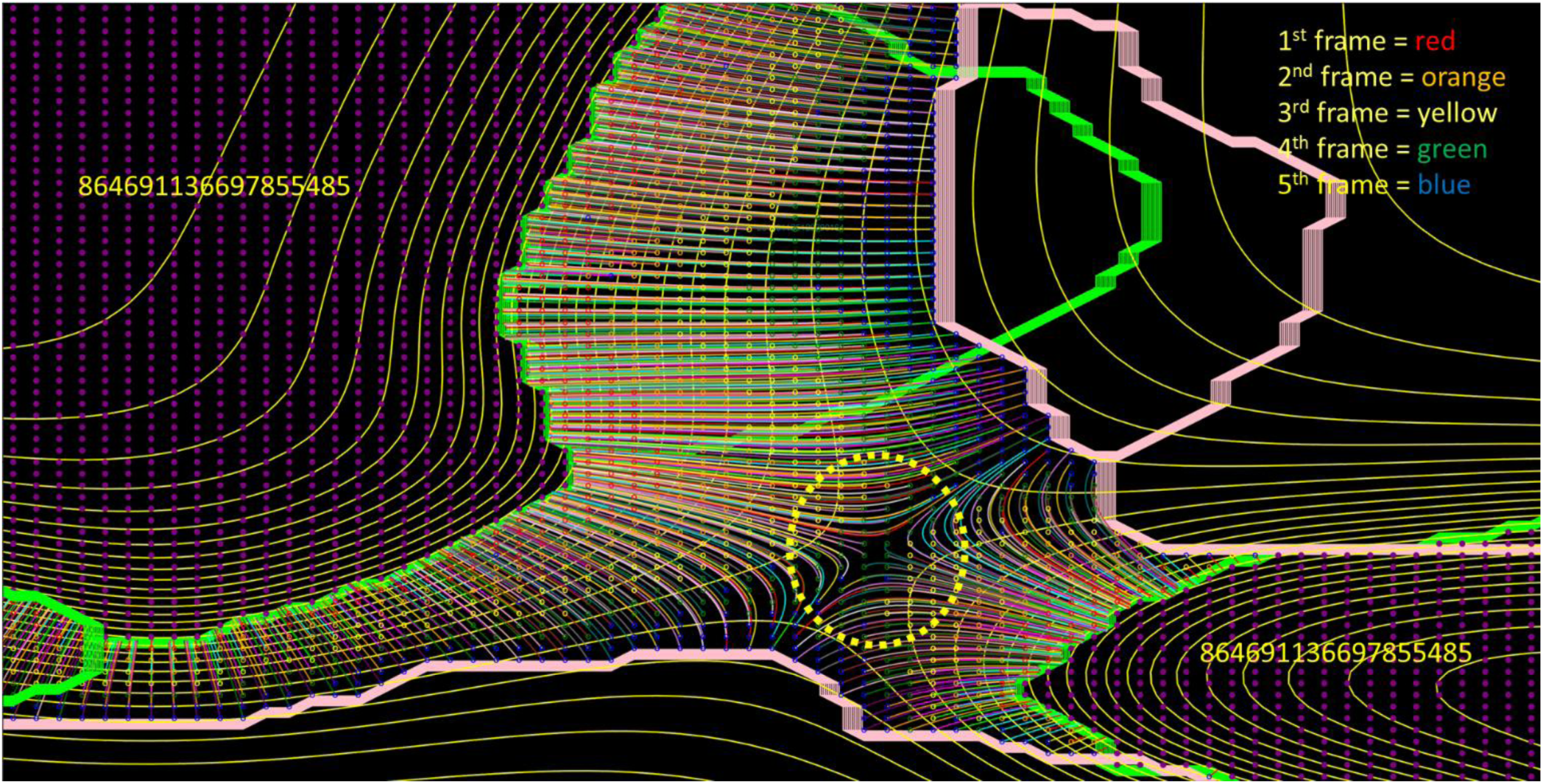
The same morphing protocol was followed as described in Figure 8, except now the two distinct regions of cell ‘485’ (upper left and lower right) fuse between the lower and upper frames. The new heat method results in a local minimum in temperature (dotted yellow circle), which leads to a smooth fusion of the cells in intermediate frames.

Pixels within the yellow circle follow the gradient to one of the overlap regions and these paths tend to be intermediate in length (color = green). Thus, the cell will fuse from two domains into one in an intermediate frame. It should be noted that identification of the green and pink boundaries is only performed for visualization. They are not utilized at all in this morphing protocol. The resulting frames are shown in Figure 12. The purple circled region shows the smooth fusion of the purple cell. The pink circled region shows a cell that appears between frames, which gradually appears and grows in size in the intermediate frames.

**Figure 12:**
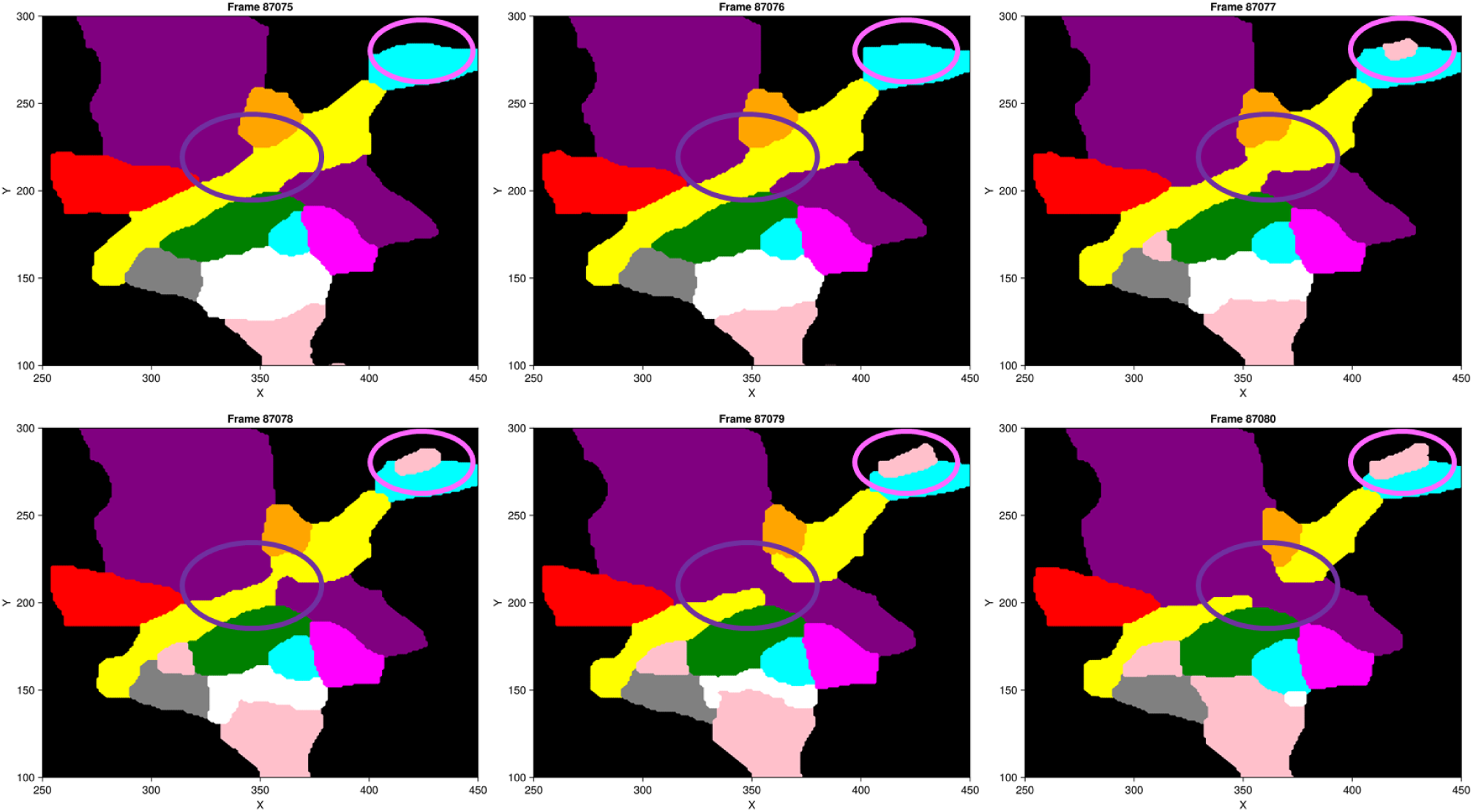
The first (zframe = 87075) and last (zframe = 87080) frames are the mode filtered segmentations. Intermediate images were generated by the heat diffusion method described in Figure 6, which was followed by additional mode filtering in the xy plane, zy plane xz plane and the xy plane again. Circled in purple, the ‘485’ cell (purple) fuses without introducing gaps or overlaps with neighboring cells. Circled in pink, part of the pink cell appears between the first and last frame and this gradual transition is shown in the intermediate frames.

Finally, the original mode filtered but stair-stepped (unmorphed) surface mesh models of this region are shown (Figure 13A) and compared with their image morphed counterparts (Figure 13B). With the purple cell removed, the correspondence between the 3D surface meshes (Figure 13C) and the 2D segmented image slice (Figure 13D) can be appreciated.

**Figure 13:**
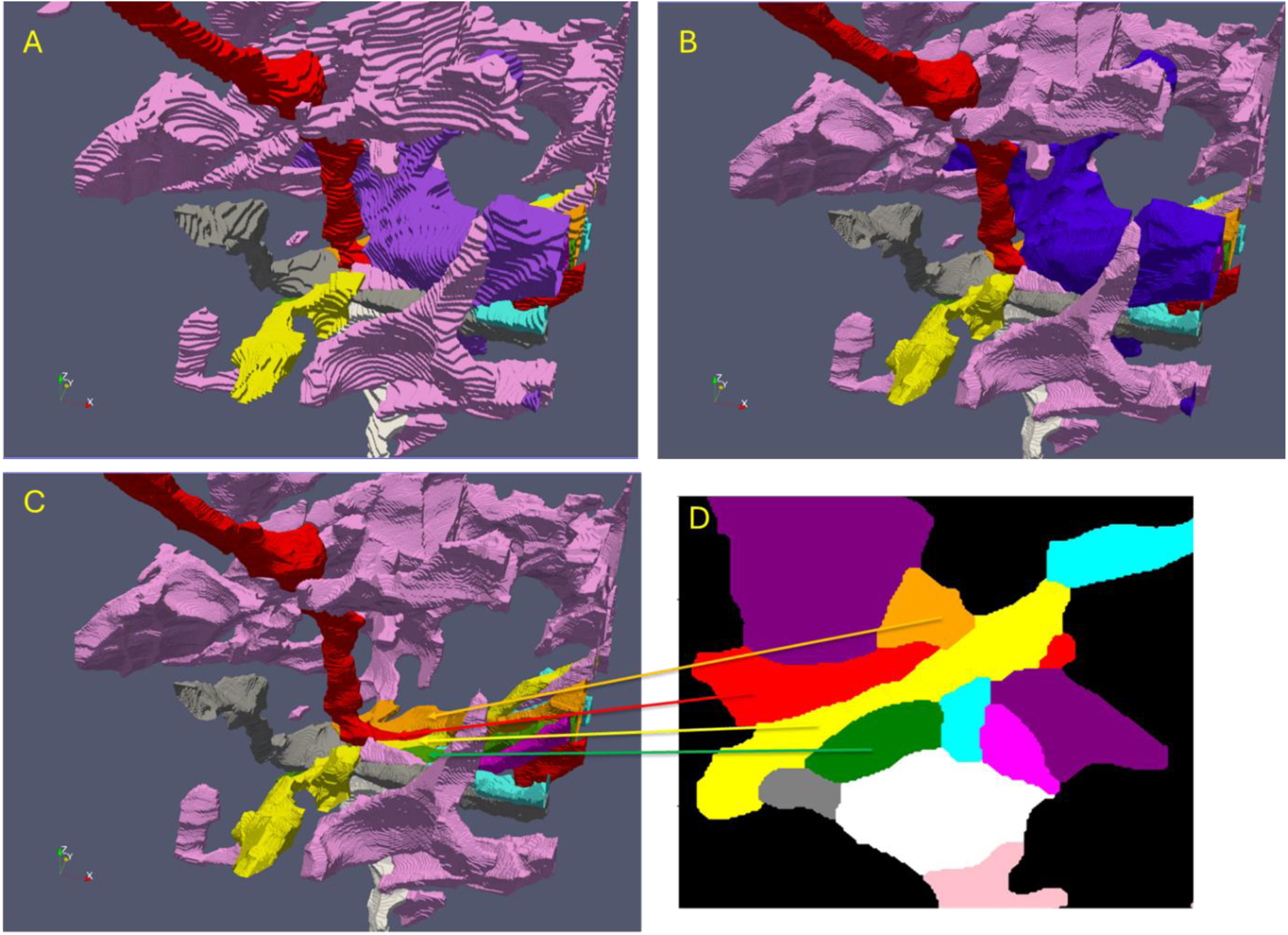
A. Stair-stepped surface meshes of the cells in the region of interest after mode filtering but before morphing. B. Surface meshes after morphing. C. Same as B, with the purple cell removed. D. Correspondence between the cells in zframe 87070 and the 3D surface meshes in panel C.

To date, the method has been applied to 2564 ‘cells’ (which dropped to 2135 ‘cells’ due to elimination of cell fragments) to produce surface meshes with high enough quality to solve the diffusion equation on the surface meshes, as presented in a companion publication.^1^ No evidence of degenerate surface meshes was found and interfaces between adjacent cells were preserved without errors.

## Discussion

A multitude of image morphing methods could be applied to the current problem. As of yet there do not appear to be metrics to judge the quality of the interpolation of TEM images of biological cells. The interpolation method should allow cells or parts of cells to smoothly appear or disappear between frames. The methods should allow parts of cells to branch or fuse between frames with an appropriate radius of curvature for biological membranes. It is possible that the branching geometries may be more complex than the examples shown here, with multiple components in both frames that must be joined in some way. The presence of isolated prominences and ledges (e.g. in Figure 2) should be avoided. Another criterion for the quality of interpolations is subtle – as shown in Figure 9, it is highly likely that the orange cell pushes the red cell down and to the left and then the orange cell is pushed back by the purple cell. Otherwise, the tendency would be to ‘pinch off’ part of the red cell as the purple cell pushes through. This is not observed here because the pinched off domain was small enough to be removed by mode filtering in zframe 87074. Thus, cells sometimes might move in coordinated ways that are not true interpolations. This is perhaps the most challenging problem and it will be referred to here as the ‘coordinated movement’ challenge. Such cases should be considered in developing metrics of frame interpolation. Another metric would quantify the smoothness of the transition, essentially how much surface area is converted in each step. This probably should be as equal as possible between frames. Developing such metrics would aid in further development of interpolation strategies.

It is important to recognize differences between the current interpolation goals and other image morphing modalities in the present application. Cross-dissolve would allow cells to bleed outside of their boundaries in the morphing process. Any leakage beyond boundaries will lead to confusing overlaps with adjacent cells or if smoothing algorithms are applied, the transition from known frame to known frame may follow arbitrary paths. The use of signed distance functions may be applicable. However, this requires identification of boundary pixels and coordination of the signed distance functions amongst the tens to hundreds of cells per image to avoid introducing gaps or overlaps between the cells.

There is another diffusion-based morphing techniques that might be applicable to the current problem with a few modifications. For example, it is possible to simply identify the regions created by the criss-crossing of cell boundaries between frames, mesh them, then apply a fixed concentration boundary condition to the overlap region and a no-flux boundary condition to the rest. Solution of the 2D heat equation should give similar results to those presented here.

However, the irregularity of the shapes of the regions may create somewhat of a meshing challenge prior to solving the heat equation. This seems like a plausible method, so a direct comparison of the methods would be interesting. Both methods likely suffer equally from the ‘coordinated movement’ problem. There may exist strategies that address the coordinated movement problem while also meeting the other criteria.

There is also the possibility that Taubin smoothing algorithms could be directly applied to stair- stepped meshes.^17^ It seems likely that the quality of some elements would suffer due to the presence of many perpendicular faces in the stair-stepped meshes, producing very small mesh elements in those regions.

An unresolved issue with the current method is its behavior near image edges. The overlap between adjacent images is only 1 pixel wide. During image morphing, the cell paths on either side of the boundary region can be radically different. Currently, the first row or column of the image of greater x or y overwrites the last row or column of the neighboring image. This leads to some stair-stepping at interfaces, which is undesirable and in need of improvement.

## Conclusions

A method was developed for interpolating frames in TEM images of biological cells from the nervous system. The method, when combined with mode filtering, handles some difficult geometries of tens to hundreds of cells without introducing errors in the surface meshes. One advantage of the method is that cell boundaries do not need to be identified or tracked.

Improvements to the method or the development of simpler alternatives may be warranted, which would be aided by the development of quantitative metrics of image interpolation quality.

## Methods

### Image_mode_filter_7.1a.jl

Image stacks are analyzed as arrays of 4 μm x 4 μm x 4 μm ‘analysis volumes’. A settings file ‘Simulation_settings _4.0_smoothed.csv’ is read with the following table columns: Simulation number, xloc (x location of the center of the base analysis volume), yloc (y location of the center of the base analysis volume), zloc (starting frame z location), height, minus_x (number of analysis volumes in the -x direction), plus_x (number of analysis volumes in the +x direction), minus_y (number of analysis volumes in the -y direction), plus_y (number of analysis volumes in the +y direction), minus_z (number of analysis volumes in the -z direction), plus_z (number of analysis volumes in the +z direction), filled (0 if z frames have not yet been interpolated, 1 following interpolation). The simulation number is given and the corresponding row records the information about the array of analysis volumes to be analyzed. Each analysis volume has a size determined by the ‘height’ parameter, which is the height or width of each image in microns. In this example, the height is 4 μm, resulting in an image size of 501 x 501 pixels with a pixel size of 8 nm. The one extra pixel in each dimension are the overlap with the neighboring images. The images are read from csv files, with each image assigned to a worker core using a job scheduler with MPI.jl.^18^ If the z-filled parameter is 0, then the images are mode filtered in the xy plane only. The images are read in with the filename pattern: frame_{zloc}_{xloc}_{yloc}_{height}.csv, where xloc, yloc and zloc are center x, center y and starting z locations, respectively, of the analysis volume and height is the size of the image stack in microns. The smoothed images are output to: sframe_{zloc}_{xloc}_{yloc}_{height}.csv.

### Diffusion_smoothing_step1_1.0b.jl

Each sframe image frame was read is read in sequentially. Bit masks were created for each unique cell in each frame. The function label_components from Images.jl was used to identify unconnected regions of each biological cell in each frame, producing a dictionary of the number of ‘components’ of each biological cell in a frame. The pixels in each cell component were stored in sparse arrays within dictionaries. The dictionaries were stored in the JLD2 format, a Julia implementation of the HDF5 format with file name: sparse_arrays_{frame}_{xloc}_{yloc}_{height}.jld2, where frame is the zloc of the frame.

### Diffusion_smoothing_step2_8.1a_mpi.jl

Two frames were placed 25 units apart and biological cell pixels were assigned a temperature of

10.0 while all other pixels are assigned a temperature of 0.0 (a temperature of 10.0 was chosen during development instead of 1.0 to improve readability if temperatures are plotted on isotherms). Other boundaries were assigned the no flux boundary condition. The finite volume method was applied on a uniform 501 x 501 x 25 grid. The center region had an initial temperature of 0.0. The linearized steady state equations were solved by the conjugate gradient method with preconditioning. The resulting temperatures in the interior region were set to 0.0 if less than 0.05 and then the remaining non-zero temperatures were stored in a sparse array. Of the 23 intermediate slices, slice 7 was chosen as the observation layer as this layer is closer to the lower frame and has isotherms that are more similar to the lower frame boundaries. This appears to lead to shorter paths to the boundary, although this has not yet been systematically tested. Results were stored with filename: result_dict_{frame}_{xloc}_{yloc}_{height}.jld2. This is by far the slowest step in the process but only needs to be performed once and the results may be used in a variety of ways in step 3.

### Diffusion_smoothing_step3_8.1a_mpi.jl

For each pair of frames, each cell’s components were analyzed for overlap. Pixels that were unique to the second frame were traced back to the overlap region by following the gradient of the temperature distribution from step 2 using gradient ascent with α = 1.0. At each time step, the location of the nearest pixel was tested for membership in the overlap region. If in the overlap region, the path was terminated and the starting pixel location and total path length were stored in a dictionary of dictionaries where the key is a tuple of the x and y coordinates of the terminal pixel. After all pixels that were unique to the second frame were analyzed, the maximum path length for each terminal pixel was recorded. Pixels within 20% of the maximum path length are converted to the cell number from the overlap region to produce the first interpolation frame. This is followed for 40, 60 and 80% of the maximum path length to produce the next three interpolation frames.

Each of the original frames is associated with four interpolated frames. The frame number for each original frame is multiplied by five and the four interpolated frames are assigned corresponding zframe numbers. The original and interpolated frames are stored as: zframe_{zloc}_{xloc}_{yloc}_{height}.csv, where zloc is now 5x the original zloc.

### Image_mode_filter_7.1a.jl

From the settings file ‘Simulation_settings _4.0_smoothed.csv’, a simulation number is specified with the z_filled parameter set to 1. In this case, the input files are of the form: zframe_{zloc}_{xloc}_{yloc}_{height}.csv, where zloc is now 5x the original zloc. Because the z_filled parameter is 1, the images are smoothed in the xy plane, the yz plane, the zx plane and the xy plane again. The ‘images’ in the other planes are generated from the entire stack of 501 x 501 x 501 pixel images. The images are output as: szframe_{zloc}_{xloc}_{yloc}_{height}.csv, where zloc is still 5x the original zloc. This is the file that is then used to produce surface meshes, as described in the companion publications.^1,2^

## Acknowledgements

The work was supported by startup funds at the University of Washington

## Author contributions

DLE designed the study, designed the framework, wrote code and wrote the paper

## Data availability

Programs used to generate the results are available at: https://github.com/elbert5770/Diffusion_morphing_v1_archive.

The source data was downloaded from the Microns Explorer website: https://www.microns-explorer.org/cortical-mm3

